# The key role of cheaters in the persistence of cooperation

**DOI:** 10.1101/2024.10.30.621164

**Authors:** Sanasar G. Babajanyan, Yuri I. Wolf, Eugene V. Koonin, Nash D. Rochman

## Abstract

Evolution of cooperation is a major, extensively studied problem in evolutionary biology. Cooperation is beneficial for a population as a whole but costly for the bearers of social traits such that cheaters enjoy a selective advantage over cooperators. Here we focus on coevolution of cooperators and cheaters in a multi-level selection framework, by modeling competition among groups composed of cooperators and cheaters. Cheaters enjoy a reproductive advantage over cooperators at the individual level, independent of the presence of cooperators in the group. Cooperators carry a social trait that provides a fitness advantage to the respective groups. In the case of absolute fitness advantage, where the survival probability of a group is independent of the composition of other groups, the survival of cooperators does not correlate with the presence of cheaters. By contrast, in the case of relative fitness advantage, where the survival probability of a group depends on the composition of all groups, the survival of cooperators positively correlates with the presence of cheaters. Increasing the strength of the social trait alone fails to ensure survival of cooperators, and the increase of the reproduction advantage of the cheaters is necessary to avoid population extinction. We validate these theoretical results with an agent-based model of a bacterial biofilm where emergence of the cooperative trait is facilitated by the presence of cheaters, leading to evolution of spatial organization. This finding contrasts the classical view that spatial organization facilitates cooperation. Our results suggest that, counterintuitively, cheaters often promote, not destabilize, evolution of cooperation.

**Significance:** Cooperation is central to the evolution of life and the emergence of multicellularity. Social traits stabilize populations against environmental volatility and enable division of labor. However, evolution of cooperation is beset by cheaters that benefit from cooperation without paying the cost. Analysis of a multilevel selection model of population evolution shows that, counterintuitively, the presence of cheaters can be beneficial or even essential for the survival of cooperators. This unexpected effect comes from multilevel selection whereby cheaters at the individual level become altruists at the group level, enabling overall growth of the population that is essential for the persistence of cooperators. These findings have broad implications for understanding the emergence of cooperation and host-parasite coevolution throughout the history of life.

## I. INTRODUCTION

Cooperation is a ubiquitous social trait, observed at every level of biological organization, spanning viruses [1, 2], bacteria [3–6], and animals [7, 8], and is essential for the emergence and survival of complex organisms and communities including human societies [9–11]. Understanding the underlying mechanisms of major transitions in evolution [12–14], such as emergence of the first cells [15–17] and multicellular organisms [18–20], requires elucidating the nature of the selective pressures that bring about cooperative (social) traits and support their persistence.

Cooperation requires individual agents to act in the interest of the community, which is not necessarily aligned with the interest of those agents themselves. Consequently, cooperative systems are vulnerable to cheaters, agents which do not contribute to but still benefit from the cooperative behavior of other group members. The emergence of cheaters imposes a relative fitness disadvantage on remaining cooperators which can eventually lead to complete loss of cooperative traits throughout the population.[21–25].

Many mechanisms supporting the emergence and persistence of cooperation, which are robust against cheating, have been theoretically described and some have been empirically characterized, including but not limited to kin selection [26–29], reciprocal interactions [28, 30–33], non-homogeneous environmental factors [34–38], indirect reciprocity [28, 39–41], and structured interaction, that is, heterogeneous interactions at the individual level in the population [42–47], and homogeneous interaction between individuals within groups in a group-structured population [48–52].

In the multilevel selection framework, which models interactions between individual agents as well as interactions between groups, potentially at multiple hierarchical levels, conflict between individual and group level selection can appear whereby a trait is disadvantageous on the individual level but advantageous on the group level, or vice versa [15, 51, 53–56]. Addressing this conflict, the emergence and persistence of social traits, that are disadvantageous at the individual level, can be enabled by restricting interactions with cheaters or by other mechanisms resulting in fitness advantage of cooperation at the group level [15–17, 48–52].

As in the case of single-level selection, in multilevel selection scenarios, the presence of cheaters is typically associated with a negative impact on the fitness of cooperators and most prior work has focused on exploring mechanisms that promote resistance to and elimination of cheaters as the only path to the survival of cooperators. Here, we demonstrate the counter-intuitive phenomenon whereby, in the context of multilevel selection, the emergence of cheaters can promote, and can even be essential, for the long-term survival of cooperators. We fully characterize a simple, general model which can be further developed to represent a wide variety of biological systems while still admitting several useful analytical results. We provide one such specific biological example in an agent-based simulation of biofilm formation for which the key model results are replicated. Further analysis and application of this framework can be expected to yield additional insights into the evolution of complex systems.

## II. RESULTS

We consider competition among groups composed of social (*A*) and asocial (*B*) individuals that differ in their reproduction rates. Asocial individuals have a reproduction advantage over social individuals at the individual level, which is independent of the presence of social individuals within the group. The last assumption differs from the setup most commonly used in evolutionary game theory, where the fitness of both social and asocial individuals is frequency-dependent at the individual level, that is, in the simplest case, is defined by a matrix game, such as the prisoner’s dilemma [8, 57]. In the context of matrix games, the fitness of a cheater in a group composed of cheaters only is lower than that of a cooperator in a homogeneous group of cooperators. In the context of the present model, the fitness of an asocial individual is independent of the composition of the group, and so, in a homogeneous group of asocial individuals, it is still greater than that of a social individual in a homogeneous group of social individuals.

Reproduction of individuals within a group leads to group reproduction via splitting, once the maximum group size *K* is reached, resulting in random allocation of individuals from the original group into each newly formed group. Individual death within a group is not considered for the sake of model simplicity, but group death is admitted. It is assumed that at this scale, social individuals provide a fitness advantage to the respective groups by decreasing the rate of group elimination. We explore both a relative and an absolute fitness advantage for cooperation to demonstrate that these two models, while yielding the same well-known birth death scenario commonly modeled throughout evolutionary biology for single-level selection [57–59], result in different fitness landscapes for the survival of social individuals in the context of multilevel selection. Incorporating the death of individuals within a group will change results quantitatively, but not qualitatively, and the same effects will be observed in a different region of the model parameter space (see SI). The environment is assumed to have a carrying capacity, *K*_*g*_, which affects group reproduction, but inter-group competition does not result in group death, that is, the death of any group in a given time step is assumed to be independent of the number of groups in the population. This major simplifying assumption is required to admit the analytical results presented. The time and group-number independent death probability reflects the limiting case of a much broader family of systems for which the total population grows at a rate which exceeds the growth rate of the subpopulation susceptible to death. The agent-based biofilm presented below illustrates one such example. The changes in group number and composition are modeled in parallel over discrete time steps during which individual reproduction in each group, group splitting, and group death may all occur, including the possibility that no event occurs. A detailed description of the above mentioned individual and group level processes is given in the Methods and Model section.

### A. Survival and growth of groups in the neutral case

To characterize the behavior of the model under conditions of competition between social and asocial individuals, we must first establish the conditions under which groups proliferate in the absence of the social trait, that is, the neutral case corresponding to *b* = 1. In this case, in a given time step,the total number of individuals in a given group *n* = *n*_*A*_ + *n*_*B*_ will increase by one with probability 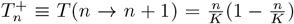 or will not change with probability 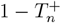 in Methods and Model). Survival and proliferation of the groups depends on the number of groups *N*_*g*_(*t*), the splitting threshold *K*, and the group elimination probability *µ* (14). Proliferation also depends on the composition of the initial groups, which in the absence of the social trait is completely determined by the number of individuals sampled uniformly from [1, *K*].

From the point of view of a given group, the survival probability at time *t* is 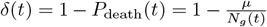, which monotonically decreases with *N*_*g*_(*t*). Given proliferation, *δ*(0) ≤ *δ*(*t*), and conversely given extinction, *δ*(0) ≥ *δ*(*t*), in the continuous limit, the population will proliferate if the derivative of *N*_*g*_(*t*) at *t* = 0 is positive. Thus, the behavior at *t* = 0 is predictive of the long term outcome. We proceeded to construct analytical estimates in which *N*_*g*_(0) ≡ *N*_*g*_ is substituted for *N*_*g*_(*t*), to yield 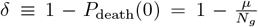 We later demonstrate that these results agree with those obtained in simulations.

Group proliferation requires that, on average, more than one daughter group survives to split again. As each group splits into two daughter groups, this requires that the probability of reaching the splitting threshold satisfies the following condition ⟨*ψ*(*δ, K*)⟩ *>* 1*/*2. This probability is found by considering the random walk on [0, *K*] averaged over the randomly chosen possible initial states *n* ∈ [1, *K*− 1] (see Methods and Model for details).

Solving ⟨*ψ*(*δ, K*)⟩ = 1*/*2 in the limit of high survival probability, *δ* ∼ 1, we find the following relation between the model parameters necessary for the proliferation of the population

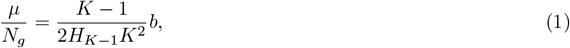

where *H*_*n*_ is the harmonic number and the reproduction scale factor *b* is also included for future reference (*b* = 1 and *b >* 1 will correspond to homogeneous groups of social and asocial individuals, respectively). From (1), it follows that for the given initial number of groups *N*_*g*_, the reproduction scale factor *b*, and the splitting size threshold *K*, there exists a corresponding threshold value of *µ* above which the population is more likely to go extinct than to reach carrying capacity. Similarly, a threshold relation exists for the number of groups *N*_*g*_ and splitting threshold *K*, for the given group death probability *µ* and the reproduction scale factor *b*. That is, there is a minimal number of initial groups *N*_*g*_ that admits the proliferation of the population with the given splitting threshold *K* for fixed *µ* and *b*. These two relations are presented in Fig.1A (black dotted lines) and in Fig.1B, respectively. Fig.1A shows the threshold relation between the splitting size, *K*, and probability of any group death *µ*, for the fixed initial number of groups *N*_*g*_ = 50 and reproduction scale factor *b* = 1. In Fig.1B, the threshold relation between *N*_*g*_ and *K* is presented for different values of *µ* and reproduction scale factor *b* = 1 and *b* = 4. The relation (1) provides the lower bounds for the model parameters, obtained in the limit of high survival probabilities *δ* ∼ 1 that allow the survival and proliferation of groups (see SI).

**FIG. 1:**
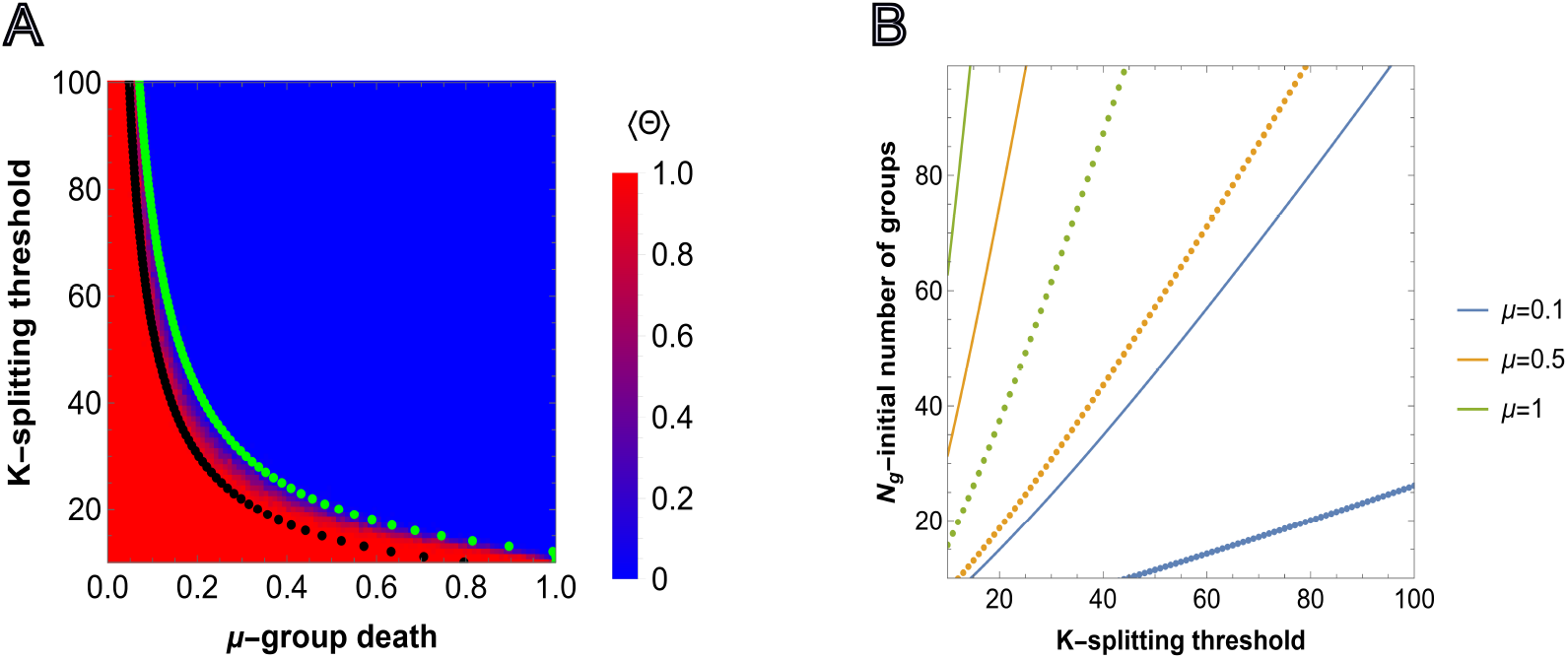
Survival of groups depending on the model parameter values. A), predictions obtained from (18) and simulation for *b* = 1 and fixed initial group size *N*_*g*_ = 50. Each cell shows the value of (2) obtained in simulations, *M* = 50 independent realizations, where the sampling is done at *T* = 1000. Steps for each pixel in the heatmap are chosen with Δ*µ* = 0.01 and Δ*K* = 1. The black and green dotted lines show the values of *µ*, for fixed *K* and *N*_*g*_, corresponding to 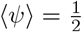 obtained under the assumption of high survival probabilities (1), and without that assumption, respectively.B), relation between the initial number of groups *N*_*g*_ and splitting threshold of a group *K* for different values of group death probability *µ* and splitting threshold *K* ∈ [10, 100] obtained from (1), below which the population goes extinct. Solid and dotted curves show the threshold values for *b* = 1 and *b* = 4 reproduction scale factors, respectively.

Complementary to (1), we consider 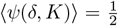 without imposing the *δ*∼ 1 condition. The green dotted line in Fig.1A shows the pairs of (*µ, K*), where *µ* is found by solving (18) as an equality for each integer value of *K* ∈ [10, 100], and for the fixed initial number of groups *N*_*g*_ = 50.

We compared the predictions from (18) with individual based simulations. For the simulations, we used an indicator function, that shows whether any group is found in the environment after time *T* or not, Θ(*N*_*g*_(*T*)) = 1 if *N*_*g*_(*T*) *>* 0, and Θ(*N*_*g*_(*T*)) = 0 otherwise.

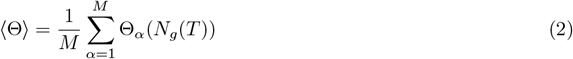

Thus, Θ = 1 means that extinction was never observed, up to time *T*, in each run of the simulation. The comparison of the simulation results and prediction of (18) is presented in Fig.1A. In Fig.SI1, the heatmaps are shown for *N*_*g*_ = 25 and *N*_*g*_ = 100.

As expected, lower splitting threshold *K* and death probability *µ*, along with larger initial number of groups *N*_*g*_, increases the probability of proliferation. Conversely, the higher the splitting threshold of groups (that is, the larger the groups), the larger the number of initial groups necessary for the population to survive and proliferate. Increasing the reproduction advantage *b* in (1) expands the region where group proliferation is possible.

The curves obtained via 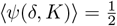 and (1) together accurately describe the results of agent-based simulation, providing upper and lower bound estimates for the model parameters that admit group proliferation.

### B. Relative fitness advantage of groups with the social trait

We assume that the social trait of *A* provides a relative advantage to groups with greater fraction of *A* by decreasing the probability of group death:

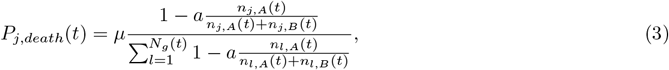

where *a* is the strength of the social trait. From (3) it follows that the survival probability of a given group depends on both the total number and the composition of all groups, thus representing density and frequency dependent selection on the group level. The social trait function was chosen to recover a well-known fitness-dependent birth-death process ([48, 57–59]), for the case when the group elimination probability is small, *µ* ≪ 1, the population reaches carrying capacity *K*_*g*_ of the environment, and the model parameters are in the region admitting proliferation of both all-*A* (*b* = 1) and all-*B* (*b* = 4) groups (see Fig.1B, further details are provided in Methods and Model section). We first consider the limit where all groups are exclusively composed of one type of individual *A* or *B* and later extend to the case of initially heterogeneous groups.

Relaxing the *µ* ≪ 1 condition, in any population of homogeneous groups, where the total number of groups *N*_*g*_ can be below the carrying capacity, the survival probabilities of all-*A* and all-*B* groups are equal to:

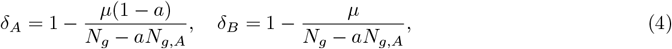

From (4) it follows that for a fixed number of groups, *N*_*g*_, the probability of any group to be eliminated from the population increases with the fraction of all-*A* groups in the population 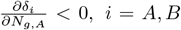. Note that this increase is not of equal magnitude between groups and *N*_*g,A*_(1 ™*δ*_*A*_) + (*N*_*g ™*_ *N*_*g,A*_)(1 ™*δ*_*B*_) = *µ* so that group death probability is independent of the composition of the population, that is with probability *µ* a group will die in a given time-step irrespective of the composition of all groups.

Indeed, for any of the all-*B* groups, the presence of all-*A* groups increases the likelihood of death due to the survival advantage of all-*A* groups provided by (3). Similarly, the presence of any other all-*A* group in the population decreases the relative advantage of each all-*A* group. The survival probabilities also depend on the social trait strength, *a*, and increasing *a* has the opposite effect on the survival probabilities of all-*A* and all-*B* groups, that is, 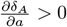 but 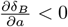.

The average probabilities of reaching the splitting thresholds for homogeneous groups 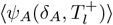 and 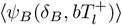 are given by (17) with their respective survival probabilities *δ*_*A*_ and *δ*_*B*_, and transition probabilities 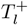 and 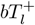 In a completely homogeneous population, that is one in which all individuals in all groups are of the same type, either all-*A* or all-*B*, the survival probability is equal for all-*A* or all-*B* cases, 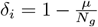, *i* = *A, B*. However, the average probability of reaching the splitting threshold for all-*B* groups is greater than that of all-*A* groups due to the reproduction advantage of *B* individuals (12,13). As a result, a homogeneous population of all-*B* groups can proliferate in some environments where a population consisting of all-*A* groups cannot (Fig.1B).

### C. Exploitation of asocial individuals in the case of relative fitness advantage

Competition among social and asocial individuals results in three possible long term outcomes: homogeneous populations of social or asocial individuals and population extinction. When the environmental conditions (the initial number of groups *N*_*g*_, splitting threshold *K*, and group death probability *µ*) are favorable for the proliferation of homogeneous populations of, the slower-growing, all-*A* groups, extinction rarely occurs and the result of competition depends on the relative strength of the social trait and asocial reproductive advantage in a straightforward manner. The resulting homogeneous population is asocial if the initial number of asocial groups is large enough and the reproductive advantage is large enough that when the total number of groups reaches group carrying capacity, if any social individuals remain, the relative fitness advantage provided by the social trait is not strong enough to prevent the stochastic elimination of this small subpopulation.

When the environmental conditions are not favorable for the proliferation of homogeneous populations of all-*A* groups, that is the environment is harsher, we observe more interesting dynamics. Note that we still consider the environmental carrying capacity, *K*_*g*_, to exceed the survival threshold values (see Fig.1B). This constraint is relaxed in the next section. Here we also continue to consider the limit where all groups are exclusively composed of one type of individual *A* or *B* and extend to the case of initially heterogeneous groups towards the end of this section. We recall group proliferation requires that, on average, more than one daughter group survives to split again and the average probabilities of reaching the splitting thresholds for homogeneous groups 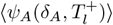 and 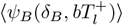 are given by (17) with their respective survival probabilities *δ*_*A*_ and *δ*_*B*_, and transition probabilities 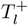 and 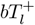. Note that ⟨*ψ*_*A,B*_⟩ depend not only on the initial number of groups in total but also the initial number of all-*A* groups and that ⟨*ψ*_*A*_⟩ is larger in the presence of all-*B* groups. Throughout the figures in the main text, we present the evaluation of initial conditions where there is an equal number of all-*A* and all-*B* groups or an equal number of social and asocial individuals, on average across simulations, in the case of heterogeneous groups.

As in the neutral case, we find the relation between the model parameters that solves 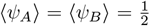, in the limit of high survival probabilities 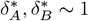. Using (20) and (4), we obtain the following relations for the threshold values of the social trait strength and the reproduction advantage of asocial individuals, *a*^*^ and *b*^*^, respectively (first (20) is evaluated for 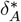 with *b* = 1, and then, for 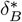 and 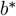, yielding the relation between *b* and *a*):

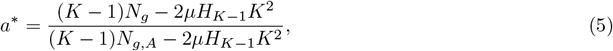

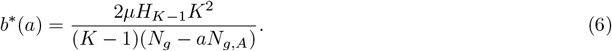

The survival probability of any group depends on the strength of the social trait but not on the asocial reproduction advantage so *a*^*^ is independent of *b* but *b*^*^ is dependent on *a*. Numerically obtained curves 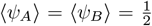 and the threshold values (5) and (6) computed in the limit of high survival probability are compared to the results from agent-based simulations in Fig.2A. The outcome of each simulation run, for various (*a, b*) pairs, is defined by

**FIG. 2:**
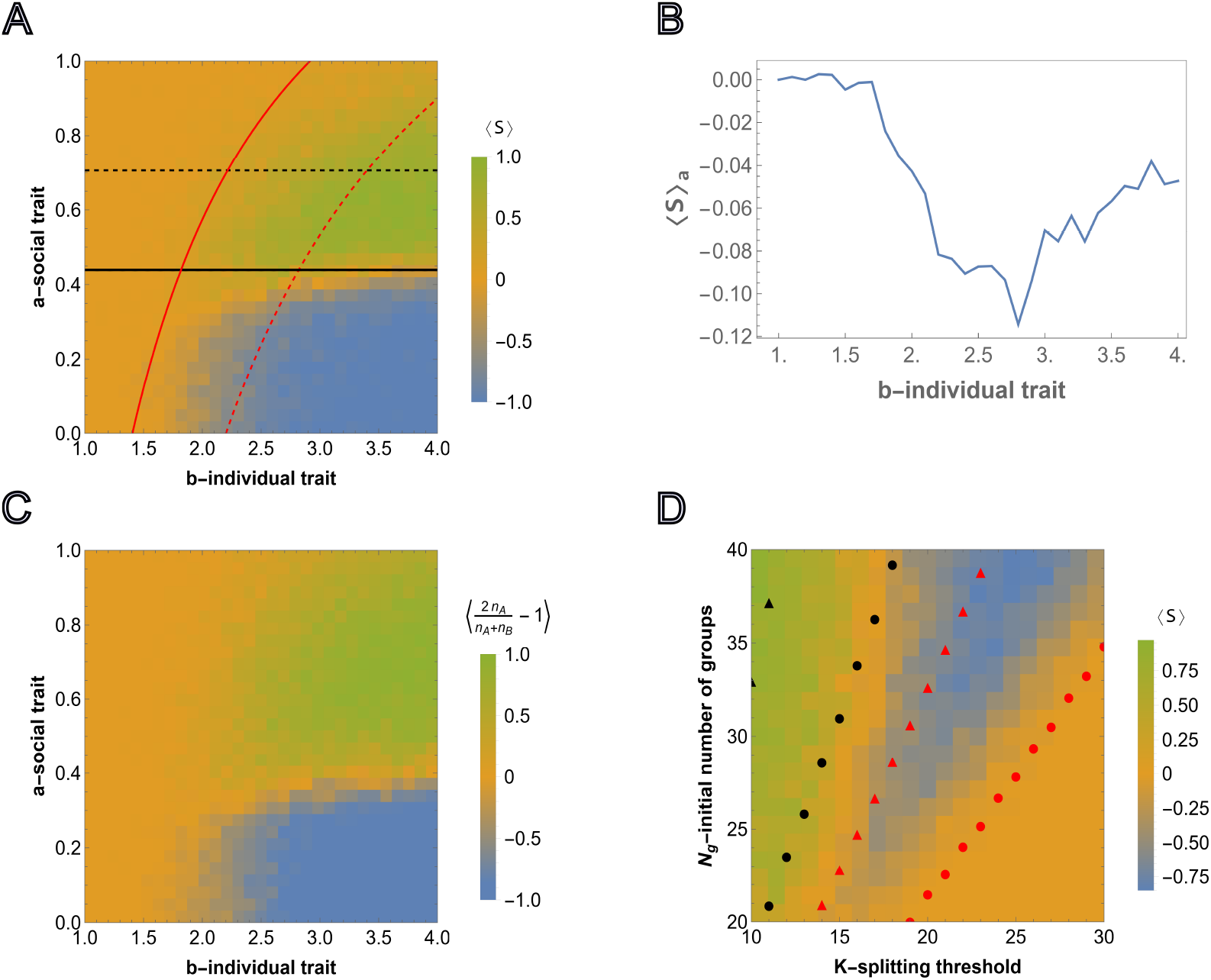
Competition between groups in the case of relative fitness advantage. A), the results of agent-based simulations are presented for the relative fitness advantage case (3)initialized with equal numbers of homogeneous groups *N*_*g,A*_ = *N*_*g,B*_. Each cell shows the value of (7) averaged over *M* = 50 independent runs, where the sampling is done at *T* = 3000. Steps for each cell in the heatmap are chosen with Δ*a* = 0.033 and Δ*b* = 0.1 starting from 0. and 1, respectively. Initial number of homogeneous groups of cooperators is *N*_*g,A*_ = 10. The black and red curves show the lines of 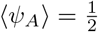 and 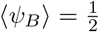, respectively. The black and red dashed curves show the threshold values of the social trait and asocial reproductive advantage, *a*^*^ and *b*^*^(*a*), respectively, such that 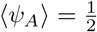 and 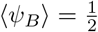. The curves are obtained from (5) and (6), respectively, B), shows the dependency of the average of ⟨*S*⟩ over all values of the social trait.,∈ *a* [0, 1] for fixed value of *b* ∈ [1, 4] C), shows the simulation results for heterogeneous intra-group composition where the initial number of each type of individuals is sampled from *U* (0, *K/*2). The heatmap shows population composition at time *T* = 5000, defined by 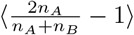, where *n*_*A*_ and *n*_*B*_ is the total number of cooperators and cheaters in the population. Extinction is assigned an output of 0. The model parameters are *µ* = 0.7, *K* = 10, *N*_*g*_ = 20 and *K*_*g*_ = 70. D), competition outcome for fixed *a* and *b* varying the group splitting threshold *K* ∈ [10, 30] and initial number of groups *N*_*g*_ ∈ [20, 40], with the initial number of all-*A* groups being equal to the nearest integer-valued lower bound (floor) *N*_*g,A*_ = [0.5*N*_*g*_], for the fixed values of social and asocial traits, *a* and *b*, respectively. The black and red circles show the values of the number of groups satisfying 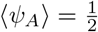 and 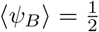 for various *K* in the considered region *N*_*g*_ ∈ [20, 40], respectively. The black and red triangles show the same 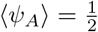 and 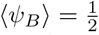, but obtained for large survival probabilities, (5) and (6), respectively. The model parameters are *µ* = 0.7, *a* = 0.4, *b* = 4, *K*_*g*_ = 150 and *T* = 5000.

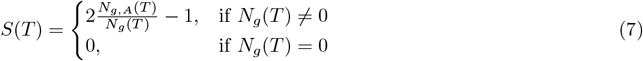

where *T* is sampling time, and *S*(*T*) = −1 and *S*(*T*) = −1 correspond to the cases when all-*A* groups out-compete all-*B* groups, respectively. The averages of *S*(*T*) are shown for *M* = 50 independent simulations 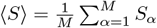 Note that ⟨*S*⟩ ∼ 0 can result either from extinction or from equal probabilities of all-*A* and all-*B* survival (neutrality with respect to the social trait, corresponding to the yellow region separating green and blue regions in Fig.2A).

Under these harsher conditions, not favorable for the proliferation of homogeneous populations of all-*A* groups, all three long-term outcomes are observed: extinction, sociality, and asociality. These regions are roughly bounded by the curves 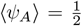 and 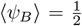. When 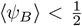 and 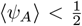, that is, all groups are more likely to die than reach the splitting threshold, the population goes extinct. When 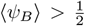 but 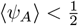, that is, all-*A* groups are more likely to die than reach the splitting threshold only all-*B* groups reach the splitting threshold and the resulting population is asocial. When 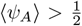 both extinction and sociality are possible. When 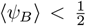, asocial groups are not able to proliferate and, consequently, the population homogenizes to all-*A* groups before carrying capacity is reached. In this case, the total number of all-*A* groups becomes too small to continue to proliferate and the population goes extinct. Extinction is prevented when 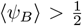. In this case, carrying capacity is reached prior to homogenization and, at carrying capacity, all-*A* groups out compete all-*B* groups resulting in a stable, social population.

This landscape illustrates the counter-intuitive dependence of the survival of the social groups on the strength of the reproduction advantage of the asocial individuals, *b*, introduced above. Indeed, ⟨*S*⟩ shows the average of simulation results for any pair (*a, b*), where *a* ∈ [0, 1] and *b* ∈ [1, 4]. Consider the average of ⟨*S*⟩ over all possible values of *a* ∈ [0, 1] for a fixed *b* (averaging over rows for single column of the heatmap matrix), that is 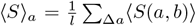 where *l* is the number of points obtained by dividing the interval [0, 1] in Δ*a* steps, shown in Fig.2B. For smaller values of *b*, ⟨ *S*⟩ _*a*_ ≈ 0, which corresponds to extinction of all groups. *S* _*a*_ *<* 0 means that for any fixed *b*, the intervals of social trait *a* where *A* wins over *B* are smaller compared to the regions where *B* wins over *A. A* wins over *B* only within the interval of *a* ∈ [0, 1], and consequently, ⟨ *S*⟩ _*a*_ is always near or below 0. ⟨ *S*⟩ _*a*_ varies nonmonotonically with respect to *b*, declining until the range within which both *B* and *A* groups survive, at which point it begins to increase again as stronger social traits are increasingly supported by the stronger reproduction advantage of asocial groups.

From this group-selection perspective, asocial *B* individuals behave as altruists. Indeed, *B* groups “sacrifice” their social-trait to reproduce quickly during the early phase of frequent group elimination so that the threshold group number admitting the fixation of *A* groups can be reached, whereas *B* groups eventually go extinct. It follows that, again counter-intuitively, the survival and proliferation of all-*A* groups is more likely when the initial fraction of these groups is smaller (see Fig.SI2a for the results with a smaller number of all-*A* groups, and Fig.SI2b for the results with the same number of all-*A* but a higher splitting threshold *K*). Conversely, from the individual-selection perspective, social *A* individuals behave as altruists “sacrificing” their reproductive advantage to the benefit of the group and asocial *B* individuals behave as “cheaters”. From either perspective, the presence of asocial, noncooperative individuals is not detrimental, but rather, is essential for the persistence of social cooperation.

In Fig.2C, the agent-based simulations are generalized to include heterogeneous groups. In each group, the initial numbers of *A* and *B* individuals are sampled from *U* (0, *K/*2). As before, the color corresponds to the relative number of social and asocial individuals in the total population across all groups at time *T*, that is, we compute 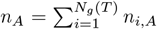 and 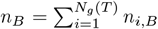 and display 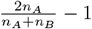 over 50 independent runs with the same model parameters (where simulations resulting in extinction are assigned the value 0). The behavior of the total population in Fig.2A and in Fig.2C is very similar indicating that the outcome at the population level is largely independent of the intra-group fixation dynamics.

We further examined the dependence of this landscape on *K* and *N*_*g*_, for fixed *a* and *b*, as shown in Fig.2D together with the analytical approximations obtained from, (5) and (6), respectively. Here, the threshold values of *N*_*g*_ and *K* are found, such that they admit proliferation of all-*A* and all-*B* groups for fixed initial fractions of all-*A* groups in the population, both in the limit of high survival probability, and as the numerically computed curves corresponding to 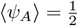 and 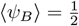. For increasing *K*, the minimum initial *N*_*g*_ required for group proliferation increases, and this threshold for cooperators is higher than that for cheaters. The numerical results approximate a lower bound for *N*_*g*_ with respect to *K* and, despite the required simplification, the analytical approximations coincide with the appropriate regions of the phase space.

### D. Propensity for sociality results in extinction in resource-limited environments

In the previous section, we demonstrated that, within a multilevel selection framework, counterintuitively, the presence of asocial, noncooperative individuals is not detrimental, but on the contrary, is essential for the persistence of social cooperation. We additionally demonstrated that when environmental conditions do not support the proliferation of a homogeneous population of social groups, but do support the proliferation of a homogeneous asocial population, social groups can outcompete asocial groups, resulting in extinction of the entire population. So far, we have assumed that the carrying capacity of the environment, *K*_*g*_, is larger than the number of groups required for the survival of a homogeneous social population (see Fig.1B) and, consequently, when *b >* 1 homogeneous asocial populations also survive. That is, for given values of *µ* and *K, K*_*g*_ is above the threshold value defined by (1) (substituting *N*_*g*_ by *K*_*g*_, and *b* = 1). Under this assumption, as long as carrying capacity is reached, extinction is prevented.

We now consider the case where *K*_*g*_ is smaller than the threshold value that supports the survival of a homogeneous population of all-*A* groups, but allows the survival of a homogeneous population of all-*B* groups, 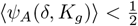 and 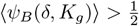, respectively (see Fig.3A). We assume that the initial number of groups *N*_*g*_ *< K*_*g*_ is such that 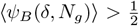. In this setting, if a sufficient number of all-A groups are present, the population will go extinct(Fig.3B, Fig.3C). Fig.3B illustrates the mixed homogeneous group case.

Here, the entire population quickly reaches the carrying capacity of the environment, primarily via proliferation of all-*B* groups. The initial proliferation of *B* groups increases the survival probability for *A* groups, leading to the subsequent decline of *B* groups, and eventually, to the collapse of the entire population. The same behavior is observed for heterogeneous group composition (Fig.3C, Fig.3D). Here, the fraction of *B* individuals in the population, 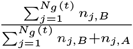, increases initially due to the reproduction advantage of *B*-dominant groups. These groups reach the splitting threshold faster than *A*-dominant groups; however, *A*-dominant groups outcompete in the long-term, eventually resulting in homogenization and subsequent extinction.

### E. Absolute fitness advantage of the social trait

In the previous sections, we assumed that sociality manifested at the group level, providing a relative fitness advantage. In this case, groups with a relatively greater *A* fraction were less likely to be eliminated than groups with a relatively smaller *A* fraction, but the overall probability of group death for a homogeneous *A* population and a homogeneous *B* population was the same. Here, we consider an alternative functional form for the social trait such that the fitness advantage provided by *A* individuals within the group is absolute, that is, independent of the composition of the population, and can affect the total probability of any group death. In this case, the probability that the given group *j* will die at time *t* is:

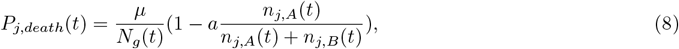

The first term in (8) is the probability that group *j* is selected at random among all *N*_*g*_(*t*) groups. The second term is the probability of death given group selection. As described above, in contrast to the previous case, Σ _*j*_ *P*_*j,death*_ ≤ *µ*. Note that *P*_*j*_ is independent of the composition of other groups, representing frequency-independence at the group level.

The function (8) results in a birth-death process with the same fixation properties as in the case of relative fitness advantage in the limit *µ* ≪ 1 (all groups are at the splitting threshold when a group dies). The survival probabilities of all-*A* and all-*B* groups are independent of population composition:

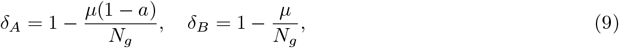

The survival probability of all-*B* groups is identical to the neutral case whereas the survival probability of all-*A* groups is rescaled with respect to the group death probability *µ* → *µ*(1 − *a*).

The threshold values of *K* and *N*_*g*_ admitting proliferation of all-*A* and all-*B* populations depend on the social trait *a* and the asocial reproduction advantage *b*. In this case, social groups outcompete asocial groups whenever 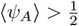, but lose otherwise. Extinction of all groups occurs when 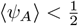 and 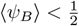 .In contrast to the relative advantage case, here, 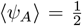 and 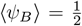 yield constant values for the social trait strength and reproduction advantage, *a* and *b*, respectively, due to the independence of the absolute fitness advantage on the population composition

The threshold values of the social trait and individual reproductive advantage strength in the limit of high survival probabilities, *a*^*^ and *b*^*^, that solve 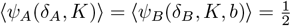 are:

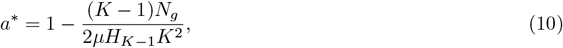

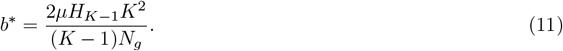

By construction, *a* ∈ [0, 1] and *b* ∈ [1, 4], and these approximations do not hold for all parameter values. That is, if the model parameters are such that *a*^*^ ∉ [0, 1] or *b*^*^ ∉ [1, 4], then the approximation in the regime of high survival probabilities does not hold. The dashed lines in Fig.4A show *a*^*^ and *b*^*^ in the limit of high survival probabilities, obtained from (10) and (11), respectively. The observed trends are qualitatively similar to those obtained with the relative advantage of the social trait (Fig.2), with a similar agreement between the analytical approximations and the simulation. In Fig.SI3, the counterparts of Fig.4 are shown, for a smaller initial fraction of homogeneous social groups, *N*_*g,A*_, and splitting thresholds *K* = 10 and *K* = 15.

We found that, when sociality confers an absolute fitness advantage that reduces the overall death probability, the persistence of social groups does not depend on the presence of asocial groups. Contrasting Fig. 4 with Fig. 3, the phase portrait is simpler than in the case of absolute fitness advantage of the social trait. Extinction is less prominent and can be prevented by increasing the strength of the social trait independent of the magnitude of the asocial reproductive advantage. Notably, in terms of the three possible long-term outcomes – sociality, asociality, and extinction – the phase portraits differ specifically with respect to extinction. By construction, both models have the same fixation properties when extinction is rare and our results for the relative fitness advantage case could not have been obtained within a framework where the number of individuals or groups is fixed (see Discussion).

**FIG. 3:**
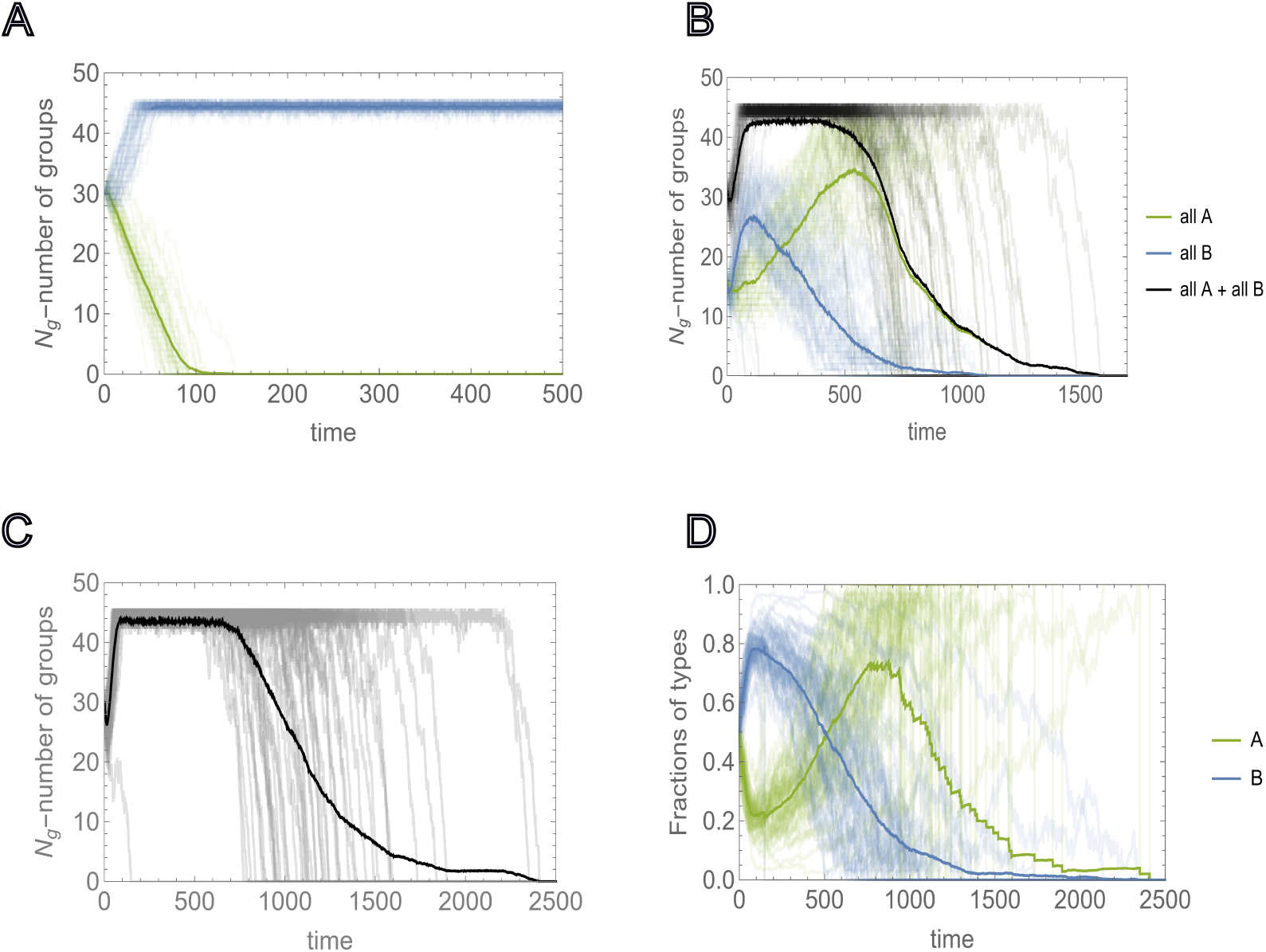
Environmental carrying capacity *K*_*g*_ and extinction due to sociality. Environmental carrying capacity *K*_*g*_ and initial number of groups *N*_*g*_ are chosen such that they allow for the proliferation of homogeneous populations of asocial groups 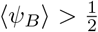, but not the proliferation of social groups, 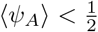 A), Extinction and proliferation of homogeneous populations of social and asocial groups (green and blue curves, respectively) for the same initial number of groups *N*_*g*_ = 30, obtained by *M* = 50 independent runs. B), Competition between all-*A* and all-*B* homogeneous groups initialized with equal numbers *N*_*g,A*_ = *N*_*g,B*_. Green and blue curves show the number of all-*A* and all-*B* homogeneous groups, respectively. Black curves show the time dependency of the total number of groups *N*_*g*_(*t*). Thick lines show behavior of the respective quantities averaged over *M* = 50 independent runs. C), Time dependence of the total number of groups, where the groups are initialized with heterogeneous intra-group composition. The initial number of each type of individuals is sampled from *U* (0, *K/*2). D), Behavior of the fractions of social and asocial individual in the population, *A* and *B*, respectively, during the time of each run of c). Thick lines show the averages of the respective quantities. In all cases, the values of the social trait and individual reproductive advantage, *a* and *b*, are chosen such that in the mixed population they satisfy (5) and (6), respectively. The model parameters are *µ* = 0.5, *a* = 0.75, *b* = 4,*K* = 20, *N*_*g*_ = 30 and *K*_*g*_ = 45.

**FIG. 4:**
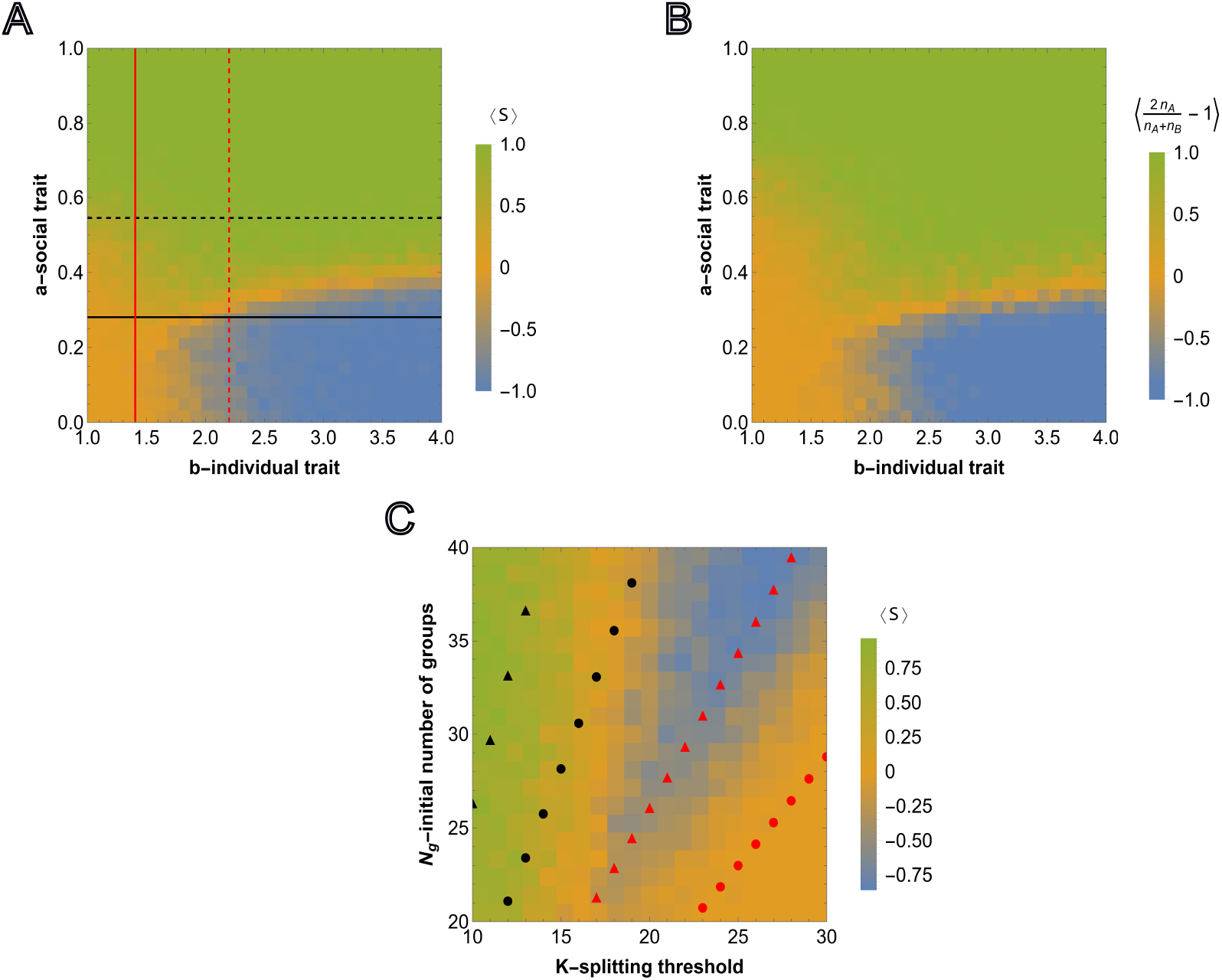
Competition outcome between social and asocial groups under absolute fitness advantage. A), Results of agent-based simulations are presented for absolute fitness advantage (8) for homogeneous groups *N*_*g,A*_ = *N*_*g,B*_. Steps for each cell in the heatmap are the same as in Fig.2A. The red and black thick lines show the values of *a*^*^ and *b*^*^, such that 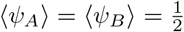. The dashed curves show *a*^*^ and *b*^*^, obtained from (10) and (11), corresponding to 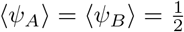 in the limit of large survival probabilities. B), Counterpart of Fig.2C is presented for the case of absolute fitness advantage. The model parameters are the same as in Fig.2A. C) Counterpart of Fig.2D for the absolute fitness advantage case with the same model parameters.

### F. Biofilm formation as an example of cheater-cooperator co-evolution

In the previous sections, we presented a general model of multilevel selection demonstrating counterintuitive dynamics in which emergence and persistence of an altruistic trait can be facilitated by or even depend on the presence of cheaters. A fundamental limitation of our approach is that, in the model, group death rate is independent of the total number of groups. More generally, these results can be expected to be qualitatively recapitulated in any system where the subset of groups that could die at any time, the “predation interface”, grows slower than the bulk population. To demonstrate an example of this behavior in a specific system, we present an agent-based simulation of biofilm growth. In our simulation, biofilms are composed of two cell types, social individuals that produce extracellular matrix (ECM) proteins which bind neighbors together[60] in exchange for a growth cost and asocial individuals which do not make ECM but can use it. Cells experience a repulsive pseudo-force acting at short distances and an attractive force at long distances. Biofilm growth is simplified to be logistic[61], constrained by an environmental carrying capacity of 500 cells. ECM production is simulated as a 100 fold increase in the attractive force constant and a 5 fold decrease in the rate of cell division. Cells are also attracted to the substrate and subject to random motion. The predation interface is composed of cells at the top of the biofilm (those which have no neighbors directly above them). Each biofilm, representing the whole population, is organized into dynamic local spatial groups. ECM production provides a relative fitness advantage to local groups of neighboring cells by decreasing the probability that cells from that group will migrate into the predation interface. It is an altruistic trait at the individual level as asocial individuals within that local group benefit from the stronger attractive pseudo-force without paying the growth cost. Consequently, in this specific system, as shown above more generally, asocial individuals are cheaters at the individual level but become altruists at the group level (see Methods for details).

At steady-state, within this parameter regime, homogeneous biofilms of ECM-producers and non-producers are approximately the same size (Fig. 5A) and, consequently, ECM production is a neutral trait at the whole-biofilm level. In contrast, in the context of colony propagation via seeding[62] by which a single individual anchors to the substrate to produce a new biofilm, ECM production is a strongly deleterious trait. The success rate for biofilm seeding is approximately 4 fold higher for the asocial non-producers (72/100 trials for non-producers;19/100 trials for ECM-producers, see Fig 5A inset). Due to the associated relative fitness advantage, mutations conferring ECM production fix with high probability within homogeneous biofilms of non-producers (99/100 trials, see Fig 5B). Fig 5C illustrates a timelapse of an example fixation simulation. It follows that the presence of cheaters facilitates the emergence and persistence of altruists in this system.

**FIG. 5:**
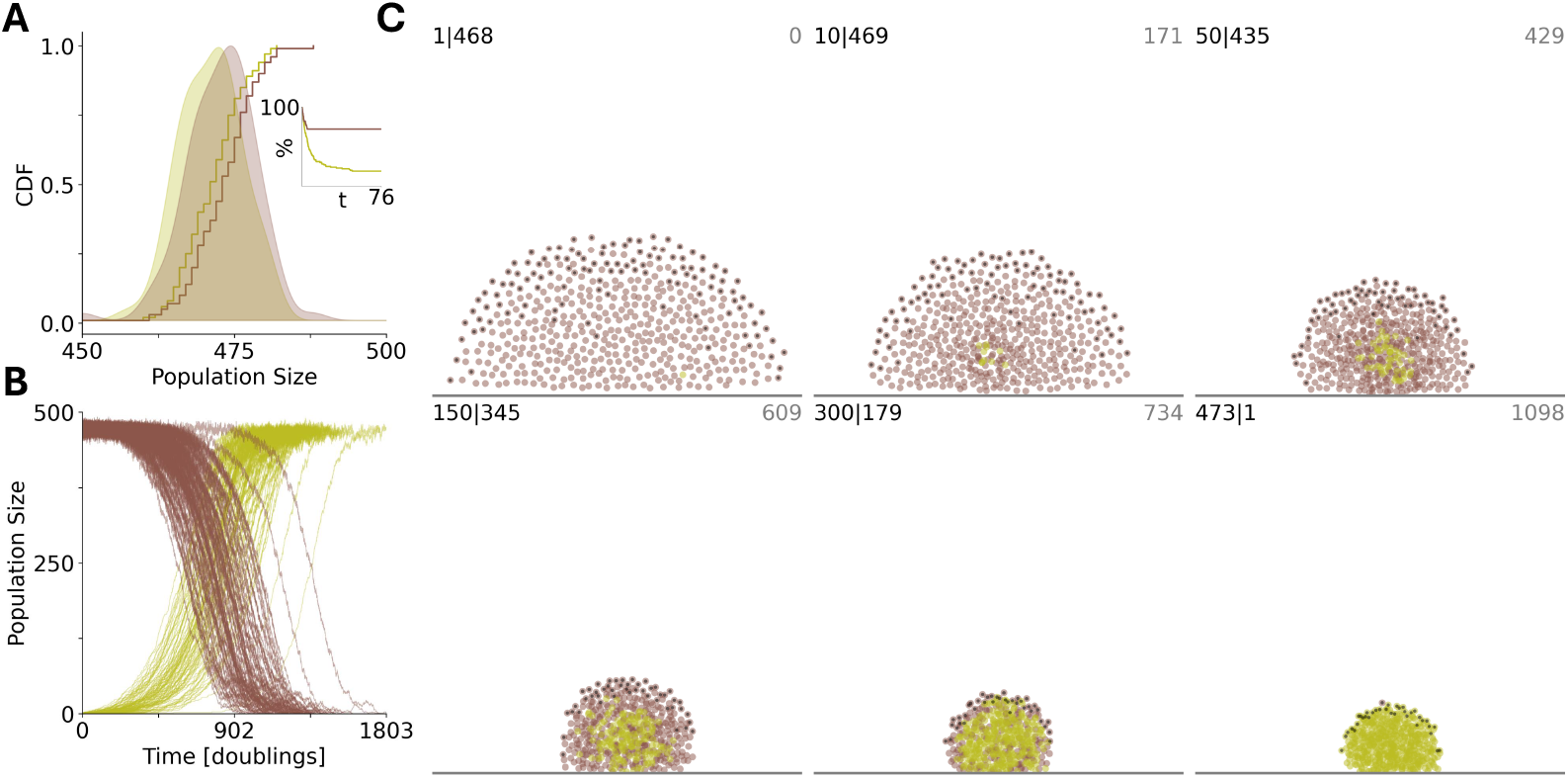
Social biofilm architects (green) benefit from asocial neighbors (brown). A), homogeneous populations are of equal size at steady-state (100 trials shown). Inset. The percentage of seeding simulations which have resisted extinction vs time. B), at time 0, a single individual acquires a mutation conferring the social phenotype in a homogeneous biofilm previously at steady-state. The mutation fixes with high probability (99 of 100 trials shown). C), timelapse of individual fixation simulation. number social— number asocial, in top left; time in top right of each panel. Individuals at the predation interface are identified in black. A/B/C. Time is measured in doubling times for asocial individuals far from carrying capacity.

## III. DISCUSSION

In this work, we consider a two-level (individual and group) selection scenario in which groups are composed of a mixture of asocial and social individuals. We demonstrate that from the individual-selection perspective, social individuals behave as altruists and asocial individuals behave as cheaters. However, although the presence of cheaters is typically associated with a negative impact on the survival and growth of social cooperators [14, 18, 57], we demonstrate the counter-intuitive phenomenon whereby, in the context of multilevel selection, the emergence of cheaters can promote and even can be essential for the long-term survival of cooperators. Our model is simple, but generalizable, and can be further developed to represent a wide variety of biological systems while still admitting several useful analytical results.

We first considered a social trait that provides a relative fitness advantage at the group level, that is, groups containing a higher proportion of individuals with the social trait (cooperators) are relatively more likely to survive than other groups. The cost of this social trait is paid at the individual level so that individuals lacking the social trait (cheaters, at the individual level) have a reproductive advantage. The reproductive advantage of the cheaters is independent of the composition of the population, as opposed to the frequency-dependent fitness more commonly assumed in evolutionary game theory [28, 48, 57].

Under these conditions, we observed three related counter-intuitive phenomena. 1) Seeding the initial population with a greater number of cheaters that do not carry the social trait tends to increase the probability that the social trait will eventually be fixed. 2) The greater the strength of the social trait, the greater the reproductive advantage of cheaters that is required to admit the proliferation of cooperator groups. 3) Cheaters can survive in environments with lower carrying capacities than cooperators thanks to their reproductive advantage; however, once carrying capacity is reached, groups of cooperators outcompete the cheaters. Consequently, the population becomes dominated by social groups, which subsequently go extinct.

The beneficial effect of asocial individuals on the survival of sociality is a consequence of multilevel selection whereby asocial individuals effectively behave as altruists at the group level. This occurs because the survival of the population – and hence the survival of social individuals – requires that the total number of groups exceeds a certain threshold, and that critical population size is maintained due to the fast proliferation of asocial individuals.

We then considered an alternative model of a social trait conferring an absolute fitness advantage. In this case, the survival of a group is defined only by its composition and does not depend on the composition of other groups in the population. In this regime, proliferation of social individuals does not depend on the initial presence of asocial individuals. Exploration of this model yielded an additional methodological insight. Both models analyzed, with a relative or absolute fitness advantage operating at the group level, result in the same classical birth-death process in the limit of infrequent group death and fixed number of groups, depending only on the strength of the social trait. In other words, the outcomes of the two models differ only with respect to extinction and whenever the social trait fixes in one case, it will fix in the other, barring extinction. It follows that evaluation of many existing models of multi-level selection[48, 50, 51, 54] where the number of individuals or groups is fixed, is inadequate to identify conditions sufficient for the evolution of social traits. In both cases, we find that lowering the group splitting threshold and increasing the initial number of groups benefits social individuals where the number of individuals or groups is fixed, is inadequate to identify conditions sufficient for the evolution of social traits.

In both cases, we find that lowering the group splitting threshold and increasing the initial number of groups benefits social individuals in this competition [15, 48, 51]. In homogeneous populations of social and asocial individuals with the same initial number of groups, cooperators can survive in smaller groups due to the reproductive disadvantage that makes them less likely to reach higher splitting thresholds. Similarly, for the same splitting threshold (maximum size of the group), asocial individuals are able to proliferate with a smaller initial number of groups than social individuals. This also holds for heterogeneous populations with either relative or absolute fitness advantages provided to groups by the social individuals.

In conclusion, we propose a simple, generalizable framework to explore evolution of cooperation that admits several useful analytical results. We validate key findings with an agent-based model of a specific system, a biofilm. We demonstrate, counterintuitively, that across a broad range of conditions, the presence of cheaters is essential for the proliferation of cooperators such that introduction of stronger social traits is insufficient for cooperation to evolve. On the contrary, stronger cooperators require stronger cheaters. Conceivably, our approach can be extended to model host-parasite coevolution, potentially, yielding a better understanding of the role of parasites in the evolution of life by multilevel selection [13]. More generally, these results stem from the frustration between the selective factors operating at different levels of organization (individual and group) which seems to underpin the evolution of complexity [63, 64].

## IV. METHODS AND MODEL

### A. Individual level interactions

Each group has a fixed number of sites, *K*, which can be occupied by resources, *R*, an *A* entity or a *B* entity. Individuals compete for available resources within each group. Let us denote the number of *A* and *B* entities within a group by *n*_*A*_ and *n*_*B*_, respectively. The available resources in this group is *K ™ n*_*A*_ *™ n*_*B*_. We assume that the resources within each group are defined at the time of the formation of the group, and no resource intake takes a place till the next reproduction of the group. Group reproduction, through splitting, can occur only when *n*_*A*_ + *n*_*B*_ = *K* (that is the resources are exhausted) whereas individual reproduction within a group can only occur when *n*_*A*_ + *n*_*B*_ *< K*. Prior to splitting, within each time step, the probability of an individual reproduction event occurring is proportional to 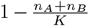, which decreases as the number of entities approaches the splitting threshold, that is, the intra-group carrying capacity. The probability that an *A* entity is the individual that reproduces is proportional to the *A* fraction of the group, 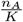 that varies over time due to the change of the number of *A* entities in the group. The relative reproductive advantage of the *B* entities, that is asocial individuals, is introduced by the scale factor, *b >* 1, yielding the transition probabilities:

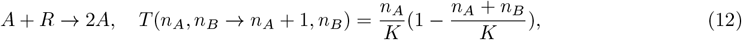

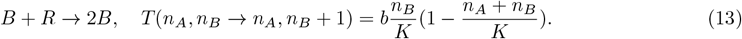

The transition probabilities must sum to no more, possibly less, than 1, that is with probability 1− *T* (*n*_*A*_, *n*_*B*_ → *n*_*A*_ + 1, *n*_*B*_) −*T* (*n*_*A*_, *n*_*B*_ → *n*_*A*_, *n*_*B*_ + 1) no updates is taking a place in the given group at the given time-step. Note, that the composition of each group is updated in parallel in a given time-step. The sum of transition probabilities is maximized when *n*_*A*_ = 0 and 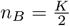 and so it follows *b* ∈ [1, 4].

### B. Group level interactions

On the group level, groups indexed over *j*, we assume that in each time step, Δ*t* = 1, a group alive at time *t* − 1 would die at time step *t* with probability:

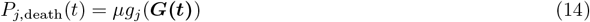

where *µ* is the probability that any group death occurs in a given time step (which is independent of group composition) and 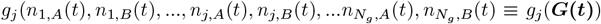 (***G*(*t*)**) which is a function of the compositions of all groups in the population, in general, reflecting the effect of the social trait. That is, the dependence of *g*_*j*_(***G*(*t*)**) on the compositions of all groups defines the relative fitness advantage of a given group compared to others. In the neutral case, that is in the absence of a social trait, 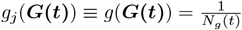 and 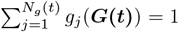. The latter condition ensures that one of the group dies with probability *µ* at any timestep. We will relax this assumption in the context of absolute fitness advantage, that is 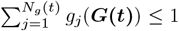 that is the probability that any group may die is less than *µ* at any time-step.

In a given time step, along with group death, which occurs with probability *µ*, group splitting occurs with probability 1 whenever at least one group has reached the splitting threshold *K* and the total number of groups is below the total environmental carrying capacity, *N*_*g*_(*t*) *< K*_*g*_. The probability that group *j* splits (a birth event) at time *t* is given by:

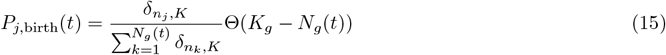

where *δ*_*k,l*_ = 1 if *k* = *l* and 0 otherwise and Θ(*x*) = 1 if *x >* 0 and 0 otherwise. If no group reached the splitting threshold *K*, then *P*_*j*,birth_(*t*) = 0 is assumed. Note that unlike group death, the probability of group splitting is independent of group composition.

Splitting of a parent group 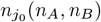 results in the formation of two daughter groups subject to the random allocation of all parent group entities yielding 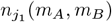 and 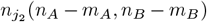 where *m*_*A*_ = *U* (0, *n*_*A*_) and *m*_*B*_ = *U* (0, *n*_*B*_) If *m*_*A*_ = *m*_*B*_ = 0 the corresponding group is immediately eliminated resulting in an abortive splitting event with probability 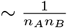.

### C. Proliferation in neutral case

For any initial state, we denote the probability of reaching the absorbing state *K* by *ψ*_*n*_. Noting that the probabilities at the boundaries *ψ*_0_ = 0, corresponding to group elimination, and *ψ*_*K*_ = 1, corresponding to the splitting threshold, are known, the remaining values of *ψ*_*n*_ may be found by solving the following recursive equation [65]

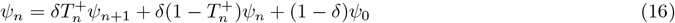

which can be understood as follows. Consider a group in state *n* at time *t*. If it survives until time *t* + 1, with probability *δ*, it can move to state *n* + 1 with probability 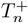 or stay in state *n* with probability 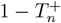 If it attains state *n* + 1, it reaches the splitting threshold with probability *ψ*_*n*+1_ (first term) and otherwise *ψ*_*n*_ (second term). The last term, corresponding to group elimination, is 0 and is included for completeness. Incorporation of individual death within groups results in a recurrence relation similar to (16) with an additional term describing transition from *n* → *n*− 1 (see SI). The analysis presented below could be carried on in that case as well, although this complicates the analysis without revealing new phenomena, but changing the regions of the model parameter space values where these phenomena are observed in the absence of individual deaths.

The solution to (16) is

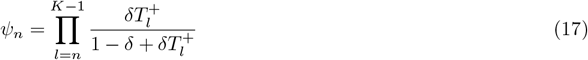

Note that, in the absence of group death, corresponding to *δ* = 1, all groups eventually reach the splitting threshold, *ψ*_*n*_ = 1. Denoting the average of (17) over all possible initial states 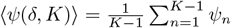, the expectation that for the given values of *µ, N*_*g*_ and *K*, groups will proliferate, satisfies the inequality:

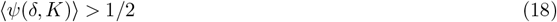

Thus, a group will proliferate if it is more likely to reach the splitting threshold than to die. The average probability of reaching the threshold increases with the increasing number of groups 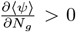, and decreases with increasing death probability 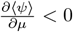 and with increasing splitting threshold value 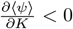. In the limit of high survival probability, *δ*∼ 1, the average probability of reaching the splitting threshold is given by (see SI for derivation):

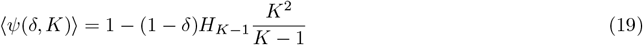

where *H*_*n*_ is the harmonic number. From (19), one can also compute the threshold value for *δ* satisfying ⟨*ψ*(*δ*^*^, *K*)⟩ = 1*/*2, that is, it is equally likely, in average, to reach the splitting threshold or die:

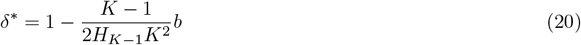

Substituting the expression for survival probability, 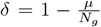, into (20), we find the relation among the model parameters (1).

### D. Relative and absolute fitness cases for *µ* ≪ 1

Under the *µ* ≪ 1 limit, let us assume that a population of homogeneous groups has reached the environmental capacity *K*_*g*_, such that *N*_*g,A*_ groups are composed of only *A* individuals and *K*_*g*_ −*N*_*g,A*_ of only *B* individuals. Indeed, for *µ* ≪ 1, all initially heterogeneous groups will homogenize first before fixation of any trait in the population. Let us further assume that each group in the population reaches the splitting threshold *K* before the next reproduction event, given that *µ* ≪ 1.This assumption also implies that group reproduction is trait-independent. Although *B* groups will, on average, reach the splitting threshold faster than *A* groups, at the carrying capacity, when group death is infrequent, all groups can be presumed to be at the splitting threshold at the time of any group death. These considerations yield the transition probabilities for relative fitness advantage case (3):

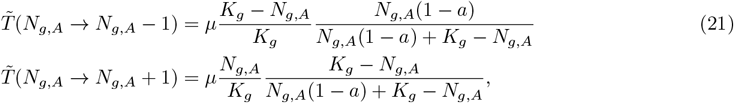

where, in the first equation, the first ratio is the probability that a *B* group splits at the time of group death and the second ratio is the probability that an *A* group was eliminated. Note that the group death probability *µ* impacts the rate of fixation but not the final state, which is determined by the ratio of the transition probabilities, 1 −*a*.

In the same limit, the absolute fitness advantage, (8), yields the following transition probabilities

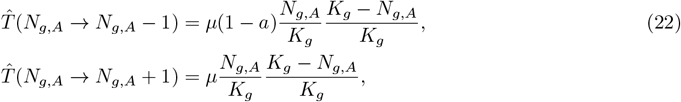

resulting in the same fixation properties dependent only on the ratio, 1 − *a*.

### E. Biofilm simulation

Biofilms reside on a substrate consisting of 400 anchor points uniformly distributed in a grid approximately twice the length of the footprint of the broadest biofilm to mitigate boundary effects. Biofilm dynamics may be subdivided into three categories: cell motion, cell division, and cell removal (death/predation). Cells move at a fixed speed of one gridpoint per timestep in a direction determined by pseudo-forces acting on them by other cells and the substrate as well as a random term. The pseudo-force between cells *p* and *q* is governed by the expression: 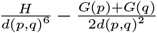 where *d*(*p, q*) is the distance separating the pair, *H* is the repulsive pseudo-force constant, and *G* is the attractive pseudo-force constant. Across all simulations shown, *H* = 1 (arbitrary) and *G* was selected to satisfy the condition: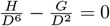, for a pair of asocial individuals, where *D* is the distance between adjacent substrate grid points. For social individuals, *G* is 100 fold larger. The pseudo-force between a single cell and each substrate anchor is of the same form where 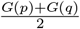 is substituted by 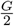 After calculating the pseudo-forces acting on each cell, the unit vector in the same direction is computed and the weighted average of it and a unit vector in a random direction is obtained: 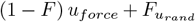 where the parameter, *F*, defines the degree to which cell motion is randomized. Across all simulations shown, *F* = 0.5. Cell positions are then updated according to movement in the specified direction and a reflective boundary condition on the substrate is imposed. Finally, a small random displacement is added to each cell position to ensure uniqueness and prevent division by zero.

After computing cell motion, cell division is implemented. Cell division depends only on the social phenotype and not on location within the biofilm or history of prior division. For each cell type, social and asocial individuals, a poisson pseudo random number is drawn with expected value: *n*_*BF*_ *r*(1 −*N*_*BF*_ */K*_*BF*_)*dt* where *n*_*BF*_ is the number of cells of the given type, *r* is the rate of cell division far from carrying capacity, *N*_*BF*_ is the total number of cells in the biofilm, *K*_*BF*_ is the carrying capacity, and *dt* is the timestep. Across all simulations shown, *r* for asocial individuals is *ln*(2) (and the doubling time far from carrying capacity is 1) and *r* for social individuals is five fold less; *K*_*BF*_ = 500; and *dt* = 0.01 (so that asocial individuals move for 100 time steps on average prior to dividing when far from carrying capacity). The selected quantity of cells of each type (or, if exceeding the total number, all cells of that type) are duplicated and a small random displacement is added to each cell position.

Following division, removal is implemented as follows. The cells farthest from the substrate along the vertical axis within any grid window are identified. The predation interface is composed of these cells. A poisson pseudo random number is drawn with expected value: *mLdt* where *m* is the number of cells within the predation interface, *L* is the rate of cell death, and *dt* is the timestep. Across all simulations shown, *L* = 0.1.

Three types of simulations were performed: the evaluation of steady-state behavior, biofilm seeding, and the fixation of a mutation conferring the social phenotype. To evaluate steady-state behavior, homogeneous populations of *K*_*B*_ cells of either type were initialized at uniformly distributed random positions within a square at the center of the substrate grid, with width 5 percent the length of the grid. Cell positions were updated for 1000 timesteps without random motion (*F* = 0), division, or removal. The full simulation then proceeded for 20000 timesteps and the final state was observed. To simulate biofilm seeding, the same procedure was followed beginning with a single cell and stopping when the population exceeded 300 cells. To simulate fixation, the biofilm was initialized according to cell positions obtained from a steady-state evaluation simulation for asocial individuals (randomly selected out of 100 trials performed). A single cell was then re-labeled as a social individual (randomly selected) and dynamics were simulated until the biofilm homogenized.

## V. ACKNOWLEDGMENTS

The authors are thankful to Joshua S. Weitz, Armen E. Allahverdyan, Christian Hilbe and Koonin group members for valuable discussions. The authors are supported through the Intramural Research Program of the National Library of Medicine, National Institutes of Health. Computations were performed using the Biowulf HPC cluster of the NIH. N.D.R. additionally received intramural support from the City University of New York Graduate School of Public Health and Health Policy.

