## Supplementary Information for "The key role of cheaters in the persistence of cooperation"

### I. INCORPORATING INDIVIDUAL DEATH INTO THE MODEL

In the main text, we do not include individual death in the model. Here, we will explore the effect of including this type of event and show that only the average probability of reaching the splitting threshold will change.

Let us consider again the neutral case whereby $b = 1$ and $a = 0$, that is, there is no difference between social and asocial individuals. Denoting the number of individual in a group by $n$, in the given time step, it is possible to have either $n \to n+1$, $n \to n-1$ and $n \to n$. To impose the normalization condition on the transition probabilities, we introduce the parameter $\nu \in [0,1]$ that controls birth vs death probabilities.

For the transition probabilities, we get

$$\hat{T}_n^+ \equiv T(n \to n+1) = \nu \frac{n}{K}(1 - \frac{n}{K}), \tag{1}$$

$$\hat{T}_n^- \equiv T(n \to n-1) = (1-\nu)\frac{n}{K}. \tag{2}$$

Note that $\hat{T}_n^+ + \hat{T}_n^- \leq 1$. The survival probability of the group $\delta$ will not change due to individual death. However, the probability to reach the splitting threshold starting from $n$, $\psi_n$, will change. In this case, the recurrence relation of $\psi_n$ has the following form

$$\psi_n = \delta \hat{T}_n^+ \psi_{n+1} + \delta \hat{T}_n^- \psi_{n-1} + \delta(1 - \hat{T}_n^+ - \hat{T}_n^-)\psi_n + (1-\delta)\psi_0 \tag{3}$$

As in the main text, the first term describes the reproduction of an individual within the group, given that the group survives. The second term describes the death of an individual given that the group survives. The third term corresponds to the case of no reproduction or death within the surviving group. The last term corresponds to the death of the group.

We proceed as described in the main text to obtain the average splitting probability in the limit of high survival probabilities and so on in general, though it would be more complicated. What follows from (3), is that the results obtained in the main text will be affected quantitatively, and will take a place in different regions of model parameters, in general. Therefore, for the sake of simplicity, we proceed by neglecting the individual death event within groups, and direct competition of the individuals within group. The last effect could be incorporated in the model in a similar way.

### II. DERIVATION OF $\langle \psi(\delta^*, K) \rangle = \frac{1}{2}$.

We start from $\psi_n = \prod_{l=n}^{K-1} \frac{\delta T_l^+}{1-\delta+\delta T_l^+}$. Observe that for $1-\delta \approx 0$ each term in the product can be written as

$$\frac{\delta T_l^+}{1-\delta+\delta T_l^+} = 1 - \frac{1}{T_l^+}(1-\delta) + O((1-\delta)^2) \equiv 1 - c_l x \tag{4}$$

where we expand the left-hand side at $\delta \approx 1$. We have denoted $c_l = \frac{1}{T_l^+}$ and $x = 1-\delta$. Then, each $\psi_n$ can be approximated as follows

$$\psi_n = \prod_{l=n}^{K-1}(1 - c_l x) \approx 1 - x\sum_{l=n}^{K-1} c_l \tag{5}$$

where we neglect the terms $O(x^2)$. The sum involved in (5) is equal to

$$\begin{aligned}\sum_{l=n}^{K-1} c_l &= \sum_{l=n}^{K-1}\frac{1}{T_l^+} = K\sum_{l=n}^{K-1}\frac{K}{l(K-l)} = K(\sum_{l=n}^{K-1}\frac{1}{l} + \sum_{l=n}^{K-1}\frac{1}{(K-l)}) = \\ &= K(H_{K-1} - H_{n-1} + H_{K-n}) = K(H_{K-1} - H_n + H_{K-n} + \frac{1}{n})\end{aligned}$$

where $H_m = \sum_{j=1}^{m}\frac{1}{j}$ is harmonic number. In the first line, we interchanged the variable $j \equiv K - l$ in the second sum. In the second line, we used the identity $H_m = H_{m-1} + \frac{1}{m}$. Now, we use the definition of the average probability to reach the splitting threshold, that is

$$\begin{aligned}\langle\psi\rangle &= \frac{1}{K-1}\sum_{n=1}^{K-1}\psi_n = \frac{1}{K-1}\sum_{n=1}^{K-1}\left(1 - xK(H_{K-1} - H_n + H_{K-n} + \frac{1}{n})\right) = \\ &= 1 - x\frac{K}{K-1}(\sum_{n=1}^{K-1} H_{K-1} + \sum_{n=1}^{K-1}\frac{1}{n}) = 1 - x\frac{K^2}{K-1}H_{K-1}.\end{aligned} \tag{6}$$

where we have used the fact that $\sum_{n=1}^{K-1} H_{K-n} - \sum_{n=1}^{K-1} H_n = 0$. Finally, from $\langle\psi\rangle = \frac{1}{2}$ and $x = 1-\delta$, we find that

$$\delta^* = 1 - \frac{K-1}{2H_{K-1}K^2} \tag{7}$$

In (4) we neglect the terms of $O((1-\delta)^2) \equiv O(x^2)$. The higher order terms in (4) are given by the following expression

$$O(x^2) = \frac{1-T_l^+}{{T_l^+}^2}x^2\sum_{i=0}^{\infty}(-1)^i(\frac{1-T_l^+}{T_l^+}x)^i \tag{8}$$

Now, (8) converges if $x < \frac{T_l^+}{1-T_l^+}$. If the last condition holds, then we get for (8)

$$O(x^2) = \frac{1-T_l^+}{{T_l^+}^2}x^2\frac{1}{1+\frac{1-T_l^+}{T_l^+}x} \tag{9}$$

which is positive and bounded again if $x < \frac{T_l^+}{1-T_l^+}$, since $T_l^+ < 1$. Note, that if $x < \frac{T_l^+}{1-T_l^+}$ holds for $l = 1$ and $l = K-1$, then it holds for all $l = 2, ..K-2$.

Let us check that $x^*$ obtained from (7) satisfy this condition. We get

$$\frac{K-1}{2H_{K-1}K^2} < \frac{K-1}{K^2}\frac{1}{1-\frac{K-1}{K^2}} \tag{10}$$

from which, we get that $x^*$ satisfies the assumption if

$$2H_{K-1} > 1 - \frac{K-1}{K^2} \tag{11}$$

which always holds as long as $K > 1$. Thus, in the (4) we neglect the positive and bounded term.

### III. SUPPLEMENTARY FIGURES

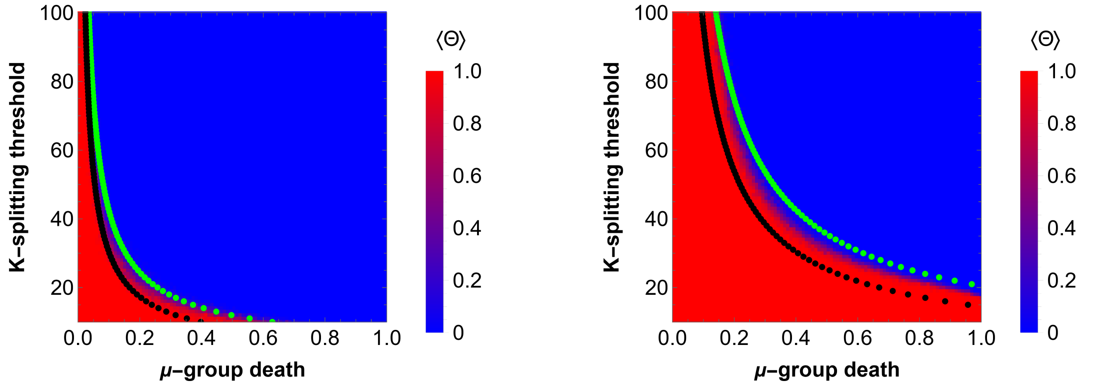


(a) (b)

FIG. 1: Survival of groups for different values of model parameters. The counterpart of Fig.1A for $N_g = 25$ and $N_g = 100$, a) and b), respectively. All the remaining parameters are the same as in the Fig.1A.

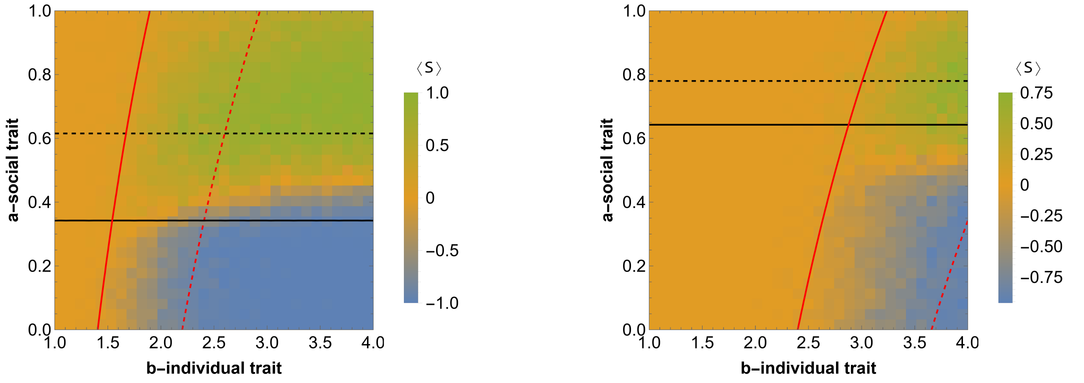


(a) (b)

FIG. 2: Competition outcome between groups of social and asocial individuals in the case of relative fitness advantage. The counterpart of Fig.2A for $N_{g,A} = \frac{N_g}{4}$, and $K = 10$ and $K = 15$, respectively. The remaining parameters are the same as in the Fig.2A.

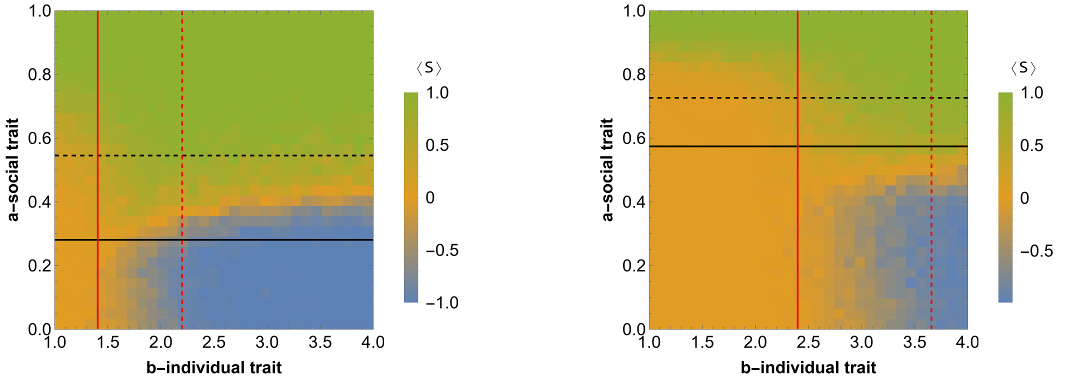


(a) (b)

FIG. 3: Competition outcome between groups of social and asocial individuals in the case of absolute fitness advantage. The counterpart of Fig.4A for $N_{g,A} = \frac{N_g}{4}$, and $K = 10$ and $K = 15$, respectively. The remaining parameters are the same as in the Fig.4A.
